# Eco-evolutionary spatial dynamics of non-linear social dilemmas

**DOI:** 10.1101/660266

**Authors:** Chaitanya S. Gokhale, Hye Jin Park

## Abstract

Spatial dynamics can promote the evolution of cooperation. While dispersal processes have been studied in simple evolutionary games, real-world social dilemmas are much more complicated. The public good, in many cases, does not increase linearly as per the investment in it. When the investment is low, for example, every additional unit of the investment may help a lot to increase the public good, but the effect vanishes as the number of investments increase. Such non-linear behaviour is the norm rather than an exception in a variety of social as well as biological systems. We take into account the non-linearity in the payoffs of the public goods game as well as the natural demographic effects of population densities. Population density has also been shown to impact the evolution of co-operation. Coupling these non-linear games and population size effect together with an explicitly defined spatial structure brings us one step closer to the complexity of real eco-evolutionary spatial systems. We show how the non-linearity in payoffs, resulting in synergy or discounting of public goods can alter the effective rate of return on the cooperative investment. Synergy or discounting in public goods accumulation affects the resulting spatial structure, not just quantitatively but in some cases, drastically changing the outcomes. In cases where a linear payoff structure would lead to extinction, synergy can support the coexistence of cooperators and defectors. The combined eco-evolutionary trajectory can thus be qualitatively different in cases on non-linear social dilemmas.

## 1 Introduction

The most significant impact of evolutionary game theory has been in the field of social evolution since a simple two player game [Axelrod, 1984] and its multiplayer version, the public goods game [Hardin, 1968] can represent the so-called *social dilemma*. The social dilemma arises when the behaviour (or choice) of an individual result in the conflict between the benefits of the individual and the group it belongs to. From decision making to biological behavioural strategies, the prisoner’s dilemma and public goods games have invited interdisciplinary studies from behavioural economists, cognitive scientists, psychologists, and biologists providing a fertile field for experimental as well as theoretical developments. While cooperative behaviour raises the group benefit, cooperators get less benefit than the others who do not cooperate arising a social dilemma. When interactions take place in a social setting where more than two individuals are involved, social dilemmas can arise in different categories. The different possible dilemmas have been categorically defined on a continuum of the so-called non-linear public goods games [Hauert et al., 2006b] as explored before by [Eshel and Motro, 1988] in the context of helping behaviour. We call the situation where the group benefit is linear in the number of cooperators *linear social dilemma*, and *non-linear social dilemma* is named after their non-linearity. Depending on the appropriate social context, it is possible that a variation of the social dilemma is more or less appropriate [Skyrms, 2003]. Archetti and Scheuring [2012] present an excellent review of the use and importance of non-linear public goods game. Interestingly, situations impossible in two player games can occur in multiplayer games which can drastically change the evolutionary outcome [Bach et al., 2006, Pacheco et al., 2009, Souza et al., 2009, Gokhale and Traulsen, 2010, Venkateswaran and Gokhale, 2018].

Of the many postulated solutions to the problem of evolution of cooperation, one of them is spatial structure. Spatial structure can be represented in different forms such as grouping, explicit space, deme structures and other ways of limiting interactions [Wright, 1930, Ohtsuki et al., 2007, Tarnita et al., 2009, 2011, Hauert and Imhof, 2012]. Especially in the repeated version of the public goods game, including an assortment mechanism promotes cooperation [van Veelen et al., 2010, 2012]. In an explicitly defined space, diffusion dynamics of cooperators and defectors support the existence of cooperators by forming spatial patterns. Comparable to the activator-inhibitor systems from the classical studies on morphogenesis by Turing [Turing, 1952], we can see various patterns with cooperators in the simplified system taking into account the linear social dilemma and constant diffusion [Wakano et al., 2009]. Previously we have combined a linear social dilemma with density-dependent diffusion coefficients [Park and Gokhale, 2019] which comes closer to analysing real movements seen across species from bacteria to humans [Okubo and Levin, 1980, Shigesada et al., 1979, Kawasaki et al., 1997, Lou and Martínez, 2009, Loe et al., 2009, Ohgiwari et al., 1992, Grauwin et al., 2009]. However, as introduced, non-linear social dilemmas have not been previously discussed in this context. Furthermore, public goods games are typically analysed in an evolutionary framework but devoid of the ecological context. Studying social dilemmas have been taken in an ecological context where along with the evolutionary change, the population dynamics are also tracked [Hauert et al., 2006a, Gokhale and Hauert, 2016, Park and Gokhale, 2019]. In this study, we aim to take the ecological context into account in non-linear social dilemmas.

In this paper, keeping the diffusion coefficient constant, we study ecological non-linear public goods games in a spatial dimension. We begin by introducing non-linearity in the payoff function of the social dilemma, including population dynamics. Then we include simple diffusion dynamics and analyse the resulting spatial patterns. For the parameter set comprising of the diffusion coefficients and the multiplication factor, we can observe the extinction, as well as heterogeneous, or homogenous patterns. Under certain simplifying assumptions, characterisation of the stability of the fixed point is possible. We discuss the dynamics of the Hopf bifurcation transition and the phase boundary between heterogeneous and homogenous patterned phases. Overall, our results suggest that synergy and discounting affects the relative size of the extinction and surviving phases. In particular, for synergy, the extinction region is reduced as the effective benefit increases resulting in an increased possibility of cooperator persistence. For discounting, the extinction region expands. The development will help contrast the results with the work of [Wakano et al., 2009] and relates our work to realistic public goods scenarios where the contributions often have a non-linear impact [Dawes et al., 1986].

## 2 Model & Results

### 2.1 Non-linear public goods game

Complexity of evolutionary games increases as we move from two-player games to multiplayer games [Gokhale and Traulsen, 2010]. A similar trend ensues as we move from linear public goods games to non-linear payoff structures [Archetti and Scheuring, 2012]. A handy method for moving from linear to non-linear multiplayer games is given in Hauert et al. [2006b]. To introduce this method in our notational form, we will first derive the payoffs in a linear setting.

In the classical version of the public goods game (PGG), the cooperators invest *c* to the common pool while the defectors contribute nothing. The value of the pool increases by a certain multiplication factor r, 1 < *r* < *N*, where *N* is the group size. The amplified returns are equally distributed to all the *N* players in the game. For such a setting the payoffs for cooperators and defectors are given by,

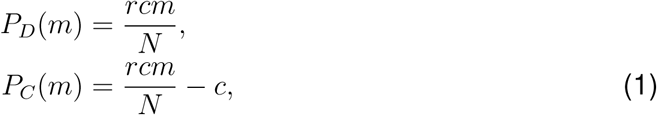

where *m* is the number of cooperators in the group. As in Hauert et al. [2006a] we are interested in not just the evolutionary dynamics (change in the frequency of cooperators over time) but the ecological dynamics as well (change in the population density over time). This system, analysed by Hauert et al. [2006a, 2008], is briefly re-introduced in our notation for later extension. We characterise the densities of cooperators and defectors in the population as *u* and *v*. Thus the population density ranges as 0 ≤ *u* + *v* ≤ 1 and the vacant space remaining in the niche is *w* = 1 − *u* − *v*. Low population density means that it is hard to encounter other individuals and accordingly hard to interact with them. Hence the group size *N*, the maximum group size in this case, is not always reachable. Instead, *S* individuals forming an interacting group. With fixed *N* the interacting group size *S* is bounded, *S* ≤ *N*, and the probability *p*(*S*; *N*) of interacting with *S* − 1 individuals is depending on the total population density *u*+*v* = 1− *w*. When we consider the focal individual, the probability *p*(*S*; *N*) of interacting with *S* − 1 individuals among a maximum group of size *N* − 1 individuals (excluding the focal individual) is,

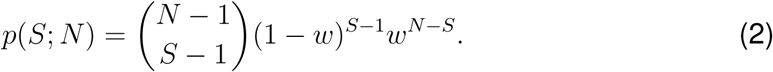

Then, the average payoffs for defectors and cooperators, *f*_*D*_ and *f*_*C*_, are given as,

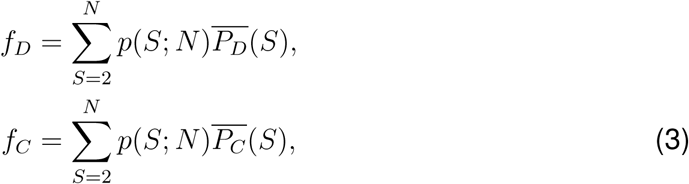

where 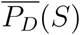 and 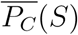 are the expected payoffs for defectors and cooperators at a given *S*. The sum for the group sizes *S* starts at two as for a social dilemma there need to be at least two interacting individuals.

To derive the expected payoffs, we first need to assess the probability of having a certain number of cooperators *m* in a group of size *S* − 1 which is given by *p*_*c*_(*m*; *S*),

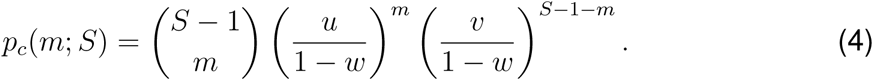

Thus the payoffs in Eq. (1) are weighted with the probability of having *m* cooperators, giving us the expected payoffs,

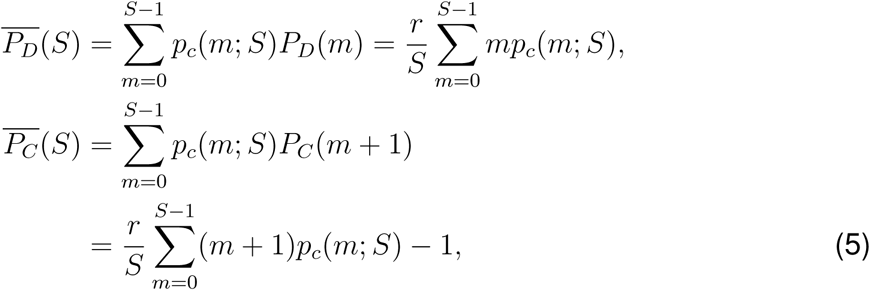

where the investment cost *c* has been set to unity without loss of generality (*c* = 1). The average payoffs *f*_*D*_ and *f*_*C*_ are thus given by,

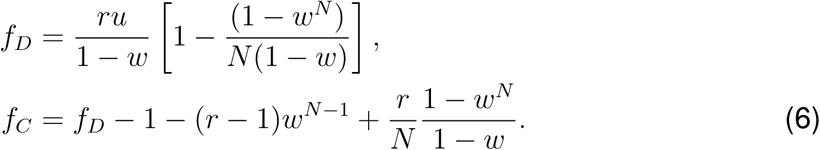

As in Hauert et al. [2006b] the parameter Ω can introduce the desired non-linearity in the payoffs as,

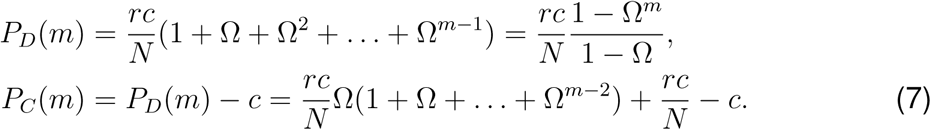

If Ω > 1, every additional cooperator contributes more than the previous, thus providing a synergistic effect. If Ω < 1, then every additional cooperator contributes less than the previous, thus saturating the benefits, thus providing a discounting effect. Following the derivation, as earlier [Gokhale and Hauert, 2016], the average payoffs are given as,

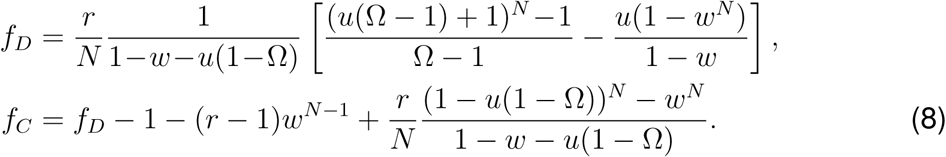

The linear version of the PGG can be recovered by setting Ω = 1.

### 2.2 Spatial non-linear public goods games

For tracing the population dynamics, we are interested in the change in the densities of cooperators and defectors over time. Both cooperators and defectors are assumed to have a baseline birth rate of *b* and death rate *d*. Growth is possible only when the population is not at carrying capacity i.e. *w* > 0. We track the densities of cooperators and defectors by an extension of the replicator dynamics [Taylor and Jonker, 1978, Hofbauer and Sigmund, 1998, Hauert et al., 2006a],

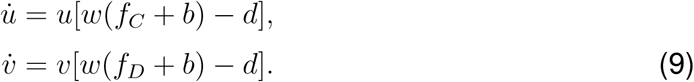

To include spatial dynamics in the above system we assume that a population of cooperators and defectors resides in a given patch. Game interactions only occur within patches, and the individuals can move adjacent patches. The patches are on a two-dimensional space and are connected in the form of a regular lattice. Taking a continuum limit, we get the differential equations with constant diffusion coefficients for cooperators *D*_*c*_ and defectors *D*_*d*_,

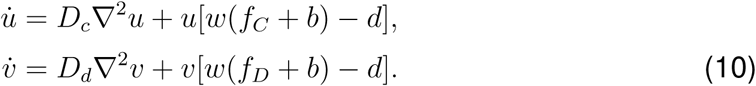

At the boundaries, there is no in- and out-flux. As in classical activator-inhibitor systems, the different ratios of the diffusion coefficient *D* = *D*_*d*_*/D*_*c*_ can generate various patterns from coexistence, extinction as well as chaos [Wakano et al., 2009].

Non-linearity in the PGG is implemented by Ω ≠ 1. Previous work shows that the introduction of Ω is enriching the dynamics [Hauert et al., 2006b, Gokhale and Hauert, 2016]. Synergy (Ω > 1) enhances cooperation while discounting (Ω < 1) suppresses it. Accordingly, synergy and discounting change the effective *r* values: With Ω larger than unity increasing *r*, and vice versa. As shown in Fig. 1, for synergy effect (Ω = 1.1), we can find a chaotic coexistence of cooperators and defectors. The same parameter for a linear case (Ω = 1.0) resulted in total extinction of the population [Wakano et al., 2009]. In the linear case, chaotic patterns were observed for *r* values larger than that of extinction patterns. Thus our observation implies the mechanism of how synergy works, by effectively increasing *r* value.

**Figure 1:**
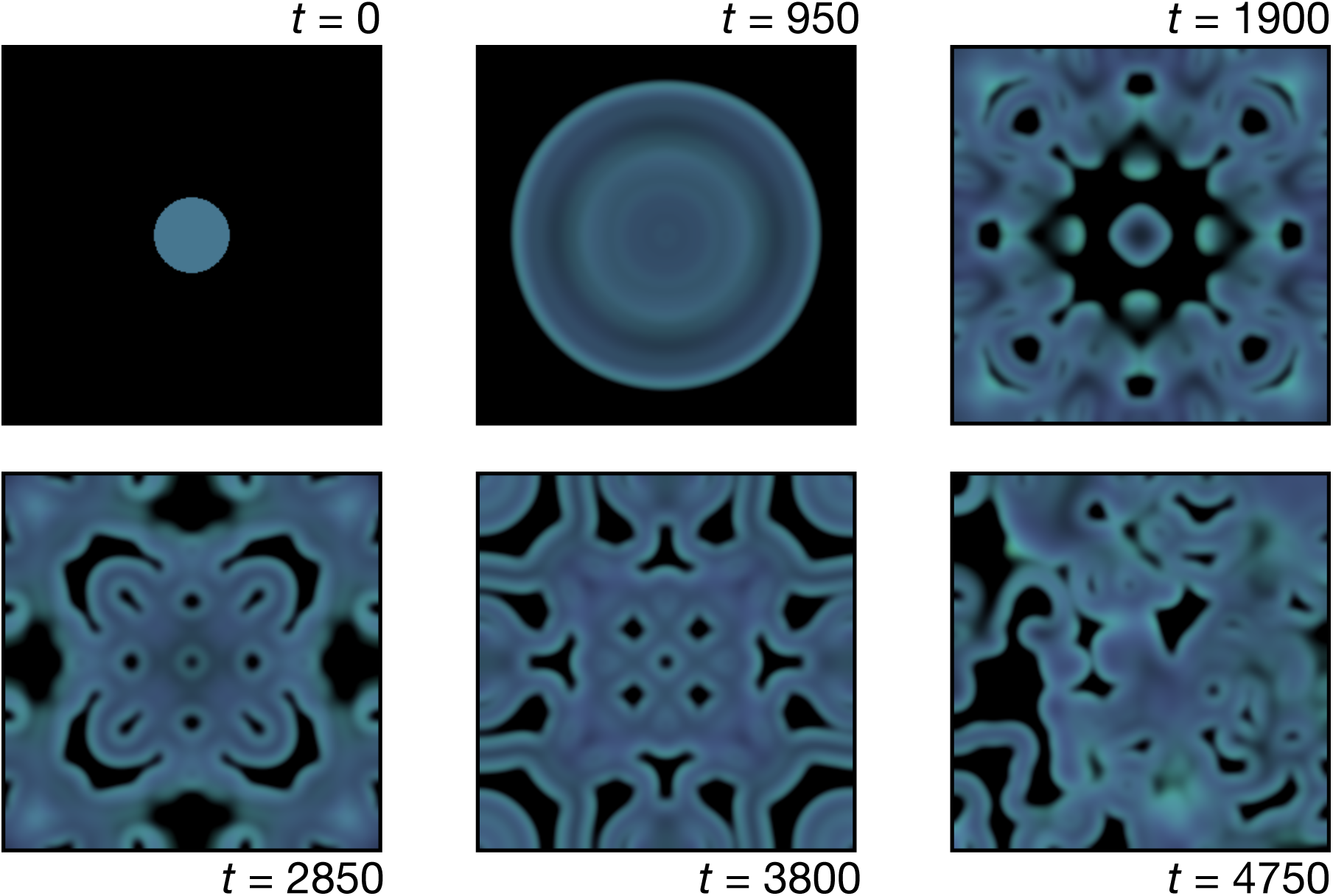
Pattern formation on the two-dimensional square lattice. We observe the chaotic pattern for Ω = 1.1 (synergy effect) where extinction comes out with Ω = 1 [Wakano et al., 2009]. Mint green and Fuchsia pink colours are used for the cooperators and defectors densities, respectively. For a full explanation of the color scheme we refer to the Appendix A. Black indicates no individual on the site whereas blue appears when the ratio of cooperators and defectors is the same. Initially, a disk with radius *L/*10 at the centre where *L* is the system size is occupied by cooperator and defector densities 0.1, respectively. We use multiplication factor *r* = 2.2 and diffusion coefficient ratio *D* = 2. Throughout the paper, for simulations, we used the system size *L* = 283, *dt* = 0.1 and *dx* = 1.4 with the Crank-Nicolson algorithm.

The change in the resulting patterns due to synergy or discounting is not limited to extinction of chaos but is a general feature of the non-linearity in payoffs. To illustrate this change we show how a stable pattern under linear PGG (Ω = 1) can change the shape under discounting or synergy in Fig. 2. Such changes in the final structure happen all over the parameter space. To confirm this tendency, we examine the spatial patterns for various parameters and find five phases, same as in the the linear PGG case [Wakano et al., 2009] but now with shifted phase boundaries (see Fig. 3). The effective *r* increases with an increasing Ω, and thus the location of the Hopf bifurcation also shifts. As a result of shifting *r*_*hopf*_, extinction region is reduced in the parameter space with synergy effect. We thus focus our attention on the Hopf bifurcation point *r*_*hopf*_.

**Figure 2:**
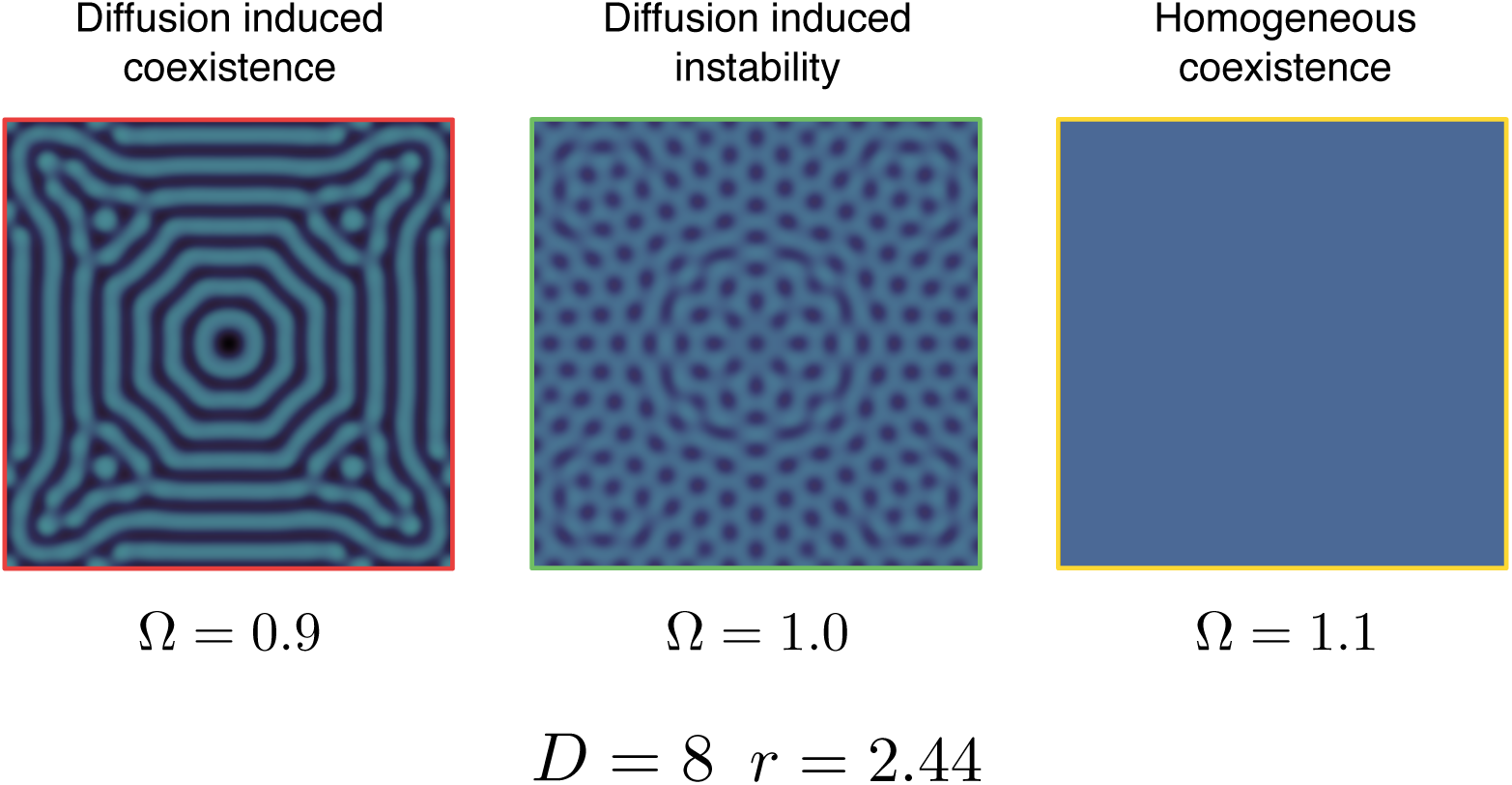
Synergy and discounting effects on pattern formation. We get the different patterns under discounting and synergy effects distinct from the linear PGG game at a given the same parameter set. While diffusion induced instability is observed in the linear PGG, the discounting effect makes diffusion induced coexistence pattern implying that the discounting effect makes the Hopf bifurcation point shift to the larger value. Under the synergy effect, on the contrary, we obtain the opposite trend observing the homogenous coexistence pattern. In the linear PGG, the homogenous patterns are observed in higher multiplication factor *r* implying the shift of *r*_*hopf*_ to the smaller value under the synergy effect. The frame colors are matched with corresponding phases explained in Fig. 3.

**Figure 3:**
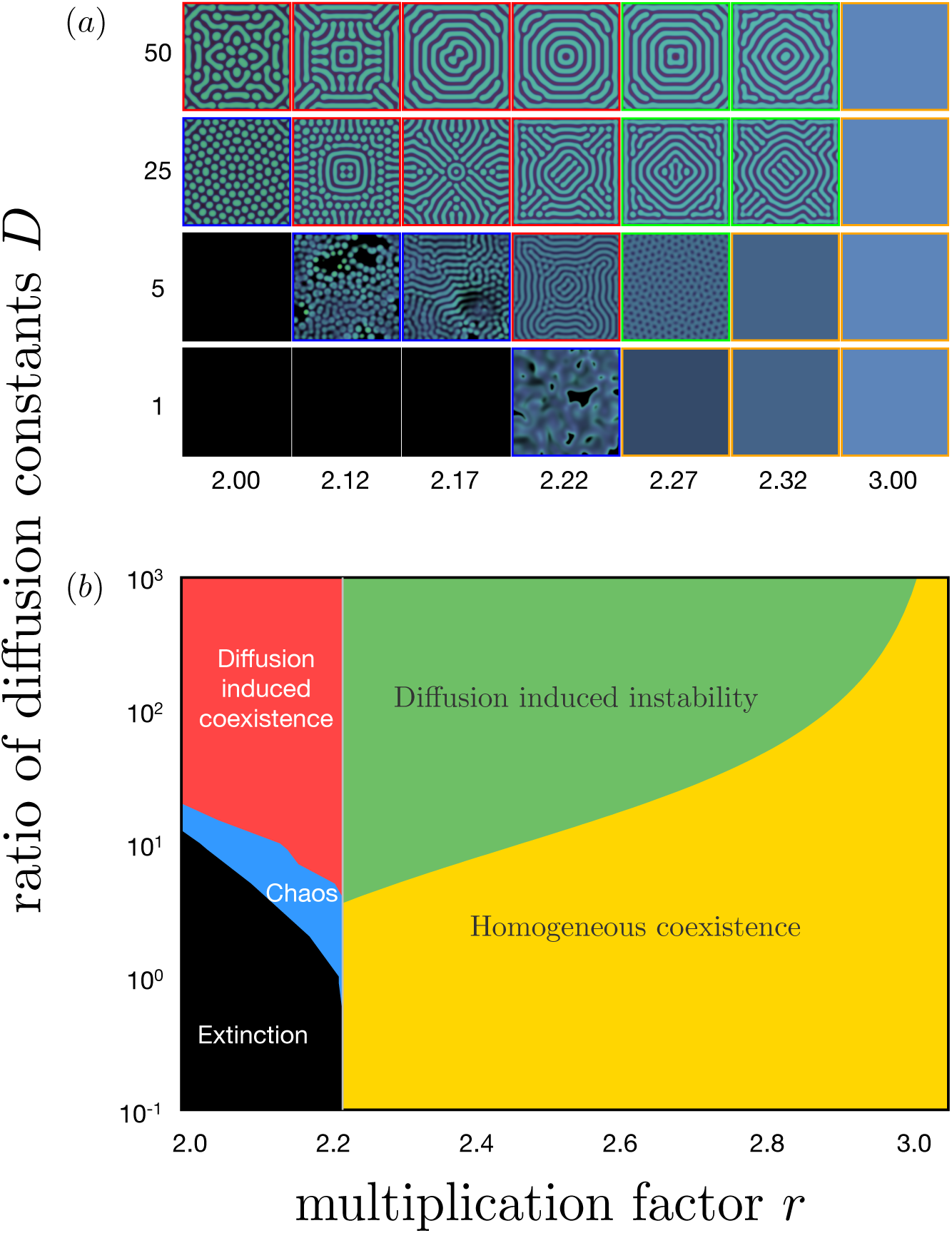
(a) Spatial patterns and (b) corresponding phase diagram for Ω = 1.1. There are five phases (framed using different colors): extinction (black), chaos (blue), diffusion induced coexistence (red), diffusion induced instability (green), and homogeneous coexistence (orange). The Hopf-bifurcation point *r*_*hopf*_ ≃ 2.2208 and the boundary between diffusion induced instability and homogeneous coexistence are analytically calculated, while the other boundaries are from the simulation results. All boundaries and *r*_*hopf*_ shift to the left indicating effective *r* increases as compared to a linear public goods game (see Fig. 4).

#### 2.2.1 Hopf-bifuraction in non-linear PGG

We find the Hopf bifurcation point *r*_*hopf*_ for various Ω values using Eq. (8). Effective *r* increases as Ω increases, and thus *r*_*hopf*_ is monotonically decreasing with Ω as in Fig 4(a). The tangential line at Ω = 1 is drawn for comparing the effects of synergy and discounting. If we focus on the differences between the tangent and *r*_*hopf*_ line, synergy changes *r*_*hopf*_ more dramatically than discounting. Synergy and discounting effects originate from 1 + (1 ± ΔΩ) + (1 ± ΔΩ)^2^ + … + (1 ± ΔΩ)^*m-*1^ in Eq. (7), where ΔΩ > 0 and plus and minus signs for synergy and discounting, respectively. Straightforwardly, the difference between 1 and (1 + ΔΩ)^*k*^ is larger than that of (1 − ΔΩ)^*k*^ for *k* > 2. Hence, the non-linear PGG itself gives different Δ*r*_*hopf*_ for the same ΔΩ.

**Figure 4:**
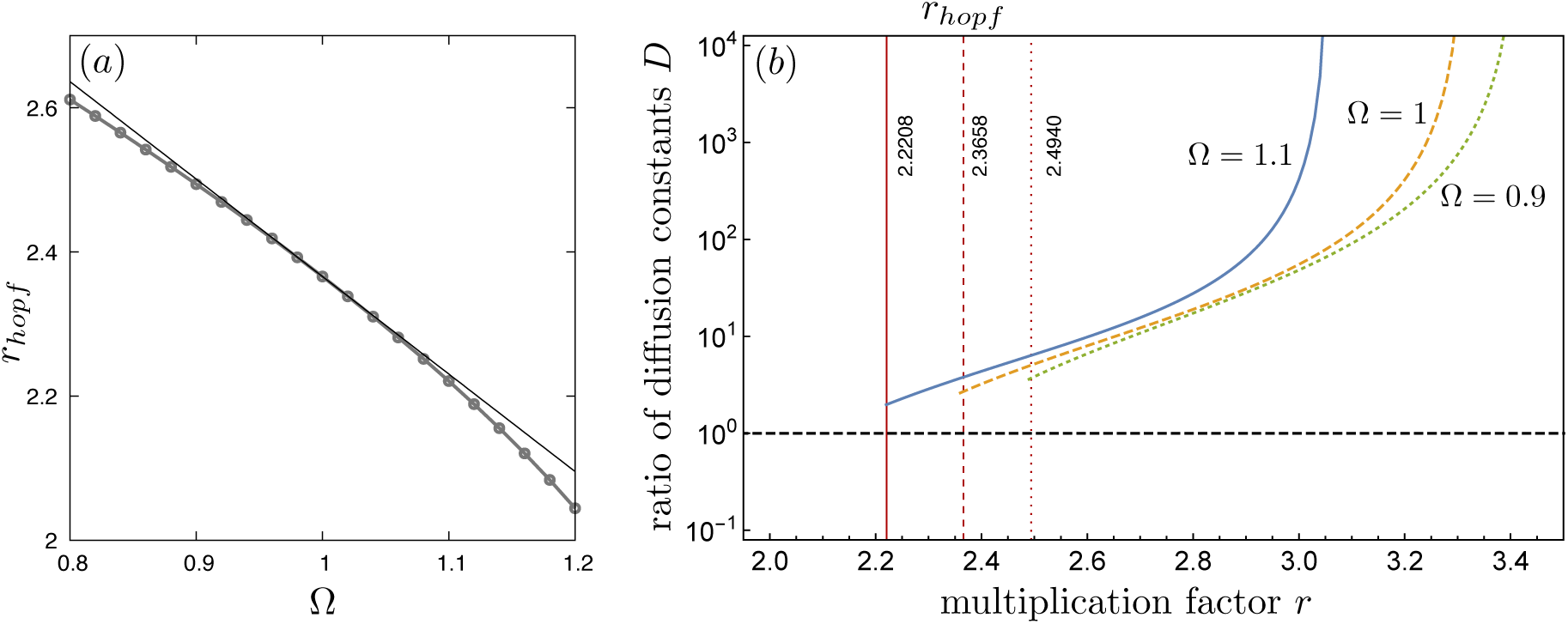
Hopf bifurcation points in Ω and shift of the phase boundary. (a) The Hopf bifurcation point *r*_*hopf*_ for various Ω (solid line with points). Synergy (Ω > 1) decreases *r*_*hopf*_ while discounting (Ω < 1) increases *r*_*hopf*_. By decreasing *r*_*hopf*_, the surviving region is extended in the parameter space. The solid line without points is a tangential line at Ω = 1. (b) The phase boundaries between diffusion induced instability and homogeneous coexistence phases are also examined for various Ω. Since *r*_*hopf*_ increases as decrease with Ω, the boundaries also move to the right.

**Figure 5:**
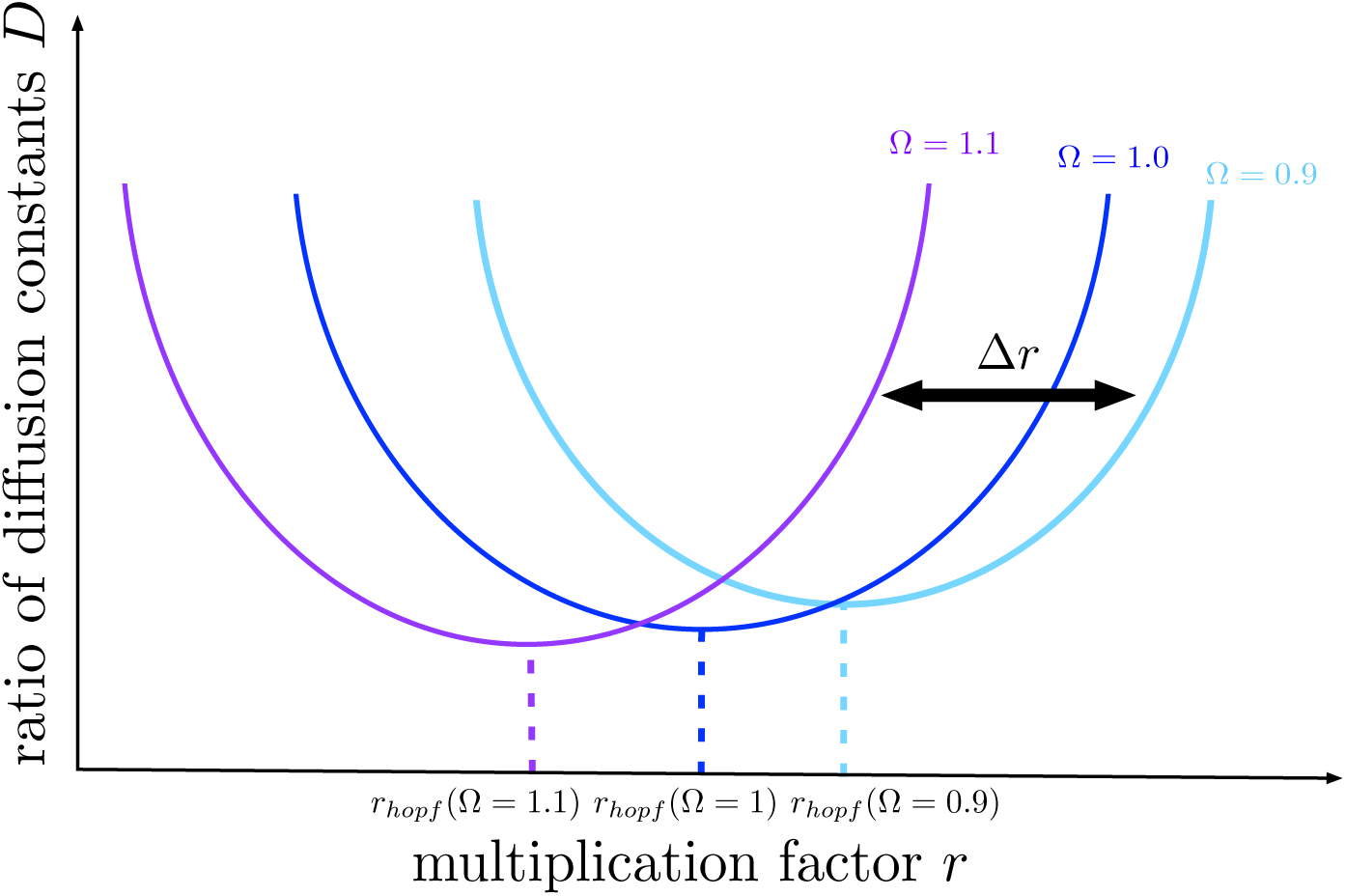
Schematic figure for expected shift of phase boundaries. According to the change of *r*_*hopf*_, over all phase boundaries may shift together at the same direction. As we have seen in Fig. 4(b), the phase boundary with *r*_*hopf*_ move to the right with discounting effect and move to the left with synergy effect, respectively. Accordingly, the surviving region in the parameter space expands with synergy effect while it shrinks with discounting effect.

#### 2.2.2 Criterion for diffusion induced instability

Since Ω changes effective *r* value, the phase boundary also moves. By using the linear stability analysis, we find phase boundaries between diffusion induced instability and homogeneous coexistence phases in *r*-*D* space shown in Fig. 4(b). To do that, we introduce new notations, and two reaction-diffusion equations in Eq. (10) can be written as

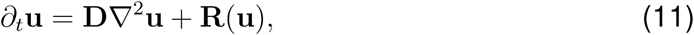

with density vector **u** = (*u, v*)^*T*^ and matrix 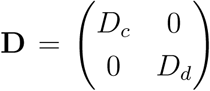. Elements of the vector 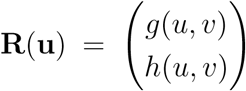 indicate reaction terms for each density which are the second terms in Eq. (10). Without diffusion, the differential equations have homogeneous solution **u**_**0**_ = (*u*_0_, *v*_0_)^*T*^ where *g*(*u*_0_, *v*_0_) = *h*(*u*_0_, *v*_0_) = 0. We assume that the solution is a fixed point, and examine its stability under diffusion.

If we consider small perturbation **ũ** from the homogeneous solution, **u** ≅ **u**_**0**_ + **ũ**, we get the relation,

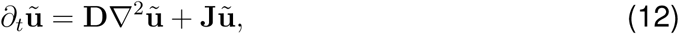

where 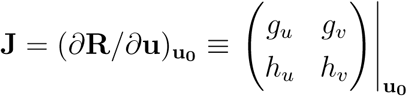. Subscripts of the *g* and *h* mean partial derivative of that variable, e.g., *g*_*u*_ means *∂g/∂u*. Decomposing 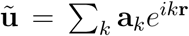 based on propagation wave number *k* gives us relation 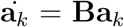 where **B** *≡* **J** − *k*^2^**D**. There-fore, the stability of the homogeneous solution can be examined by the matrix **B**. Note that Tr(**B**) < 0 is guaranteed because Tr(**J**) < 0. Hence, if the determinant of **B** is smaller than zero [det(**B**) < 0], one of the eigenvalues of the matrix **B** is positive. Then, the homogeneous solution becomes unstable and Turing patterns appear.

The condition for det(**B**) < 0 is given by

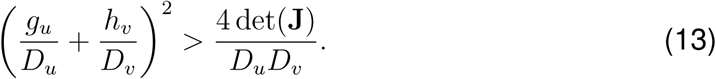

It can be rewritten as following form

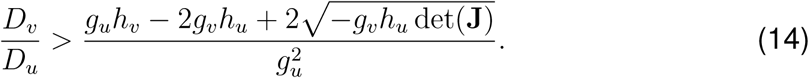

If the above criterion is satisfied, the stable fixed point predicted without diffusion becomes unstable due to diffusion. From this criterion, we get the analytic phase boundary for *r*_*hopf*_ < *r* as shown in Fig. 4(b).

## 3 Discussion

Linear public goods game is a useful approximation of the real non-linearities in applications of social dilemmas from the micro to the macro scale [Turner and Chao, 1999, Gore et al., 2009, Packer and Ruttan, 1988] with application such as in cancer [Aktipis, 2016] as well as antibiotic resistance [Lee et al., 2010]. However, when non-linearities are taken into account, the resulting outcomes might often be different from what is naively expected [Gokhale and Hauert, 2016]. In this manuscript, we have extended the analysis of spatial public goods games beyond the traditional linear public goods games. The benefits in our case are accrued in a non-linear fashion in the number of cooperators in the group. Each cooperator can provide more benefit than the last one as the number of cooperators increases (resulting in synergy) or each cooperator provides a smaller benefit than the previous one (thus leading to discounting) [Hauert et al., 2006b]. Such an extension to public goods games was proposed very early on by Eshel and Motro [1988]. Termed as superadditivity in benefits, extending from the paper one can visualise non-linearities cropping up in the costs as well, a concept not yet dealt with.

Again, such economies of scale [Dawes et al., 1986] can be justified in both bacterial as well as human interaction as proxies for quorum quenching or accruing of wealth (or austerity) [Archetti, 2009, Archetti and Scheuring, 2010, Penã et al., 2015]. Non-linearities in interactions have a profound effect when it comes to fecundity and avoiding predation be being in a group [Zöttl et al., 2013, Wrona and Jamieson Dixon, 1991]. We show that including such non-linearities in the benefit function affects the effective rate of return from the public goods game, irrespective of the types of diffusion dynamics. Just as in a non-spatial case, synergy can improve the level of cooperation in a population, in the spatial case, synergy increases the effective rate of return on the investment and expands the surviving region in the parameter space. This itself may make cooperation a favourable strategy. It would be interesting to see if the stability of the patterns is maintained as Ω switches between synergy and discounting over time [Gokhale and Hauert, 2016]. Such seasonal variations in the rate of return fundamentally change the selection pressures on cooperation and defection and can lead to not just richer evolutionary dynamics [McNamara, 2013] but eco-evolutionary spatial dynamics.

## Acknowledgements

We thank Christoph Hauert for comments and suggestions on a previous version of the study. Both authors acknowledge generous support from the Max Planck Society.

## A Colour coding

Similar to the colour coding used in Park and Gokhale [2019] we use mint green (color code: #A7FF70) and Fuchsia pink (color code: #FF8AF3) colors for denoting the cooperator and defector densities respectively for each type. The colour spectrum and saturation is determined by the ratio of cooperators to defectors which results in the Maya blue color for equal densities of cooperators and defectors. For convenience, we use HSB color space which is a cylindrical coordinate system (*r, θ, h*) = (saturation, hue, brightness). The radius of circle *r* indicates saturation or the color whereas *θ* helps us transform the RGB space to HSB. The total density of the population *ρ* = *u* + *v* is represented by the brightness *h* of the color. For better visualization, we formulate the brightness *h* as

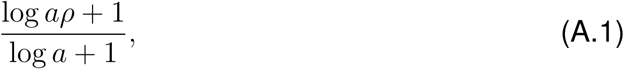

where a control parameter *a* (> −1 and ≠ 0) (see Fig A.1). The complete color scheme so developed passes the standard tests for colourblindness.

**Figure A.1:**
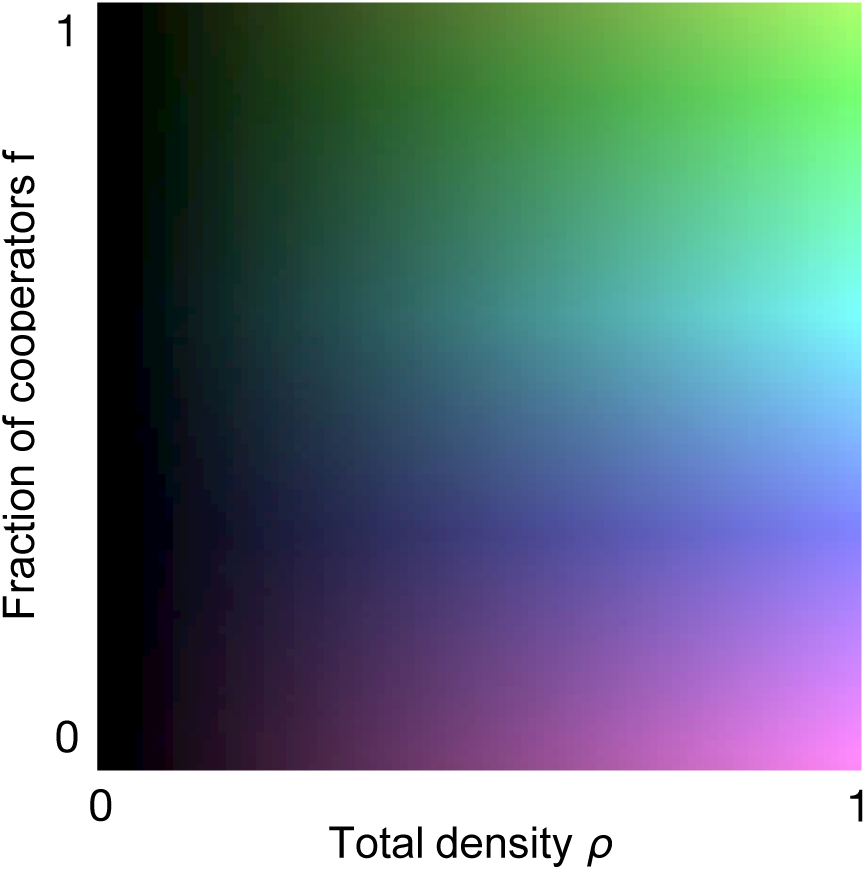

The exact color scheme developed for coloring the patterns. Each patch in a pattern is colored using this palette by choosing the corresponding *f* and *ρ* values. For brightness we used Eq. (A.1) with *a* = 15.

